# Quantification of soy-based feed ingredient entry to the United States by ocean freight shipping and the associated seaports

**DOI:** 10.1101/815134

**Authors:** Patterson Gilbert, Niederwerder Megan, Dee Scott

## Abstract

The potential of feed ingredients to serve as vehicles for African Swine Fever Virus (ASFV) introduction to the US is a significant concern. ASFV DNA has been detected in the Chinese feed system; raw grains and meals drying on the ground and milling facilities and feed delivery vehicles. Experimental evidence of ASFV survival in multiple soy-based feed ingredients during a simulated 30-day transoceanic journey and the transmission of ASFV through the natural consumption of contaminated feed has been published. Therefore, it’s important to understand the quantity of soy-based ingredients that enter the US from ASFV-positive countries via ocean shipping and rank sea ports of Entry (POEs) according to annual volume of these products to manage this risk.

The quantity of soy-based feed ingredients and their specific ports of entry was obtained at the International Trade Commission Harmonized Tariff Schedule website (www.hs.usitc.gov), a publically available website that provides a transaction of specific trade commodities between the US and its international trading partners. A close review of this database identified 10 HTS codes pertaining to soy-based feed ingredients, including soybeans, soybean meal, soy oil cake and soy oil. Specific queries on these 10 HTS codes were designed to provide information on country of origin, quantity of product, date of entry, and POE into the US. Data were exported into Microsoft Excel, then organized into pivot tables that described the quantity of specific product by country of origin and POE. The analysis focused on the 43 ASFV-positive countries on the Canadian Food Inspection Agency Watch List.

In 2018, 104,707 metric tons (MT) of soy-based ingredients were imported to the US from a total of nine foreign countries that are included on the CFIA Watch List. 52.6 % of this volume that was imported, or 55,101 MT, originated from China. These soy-based products from China entered the US from a total of 13 separate ports of entry (POEs). Of these POEs, a total of 4 POEs received greater than 88% of all of soy-based ingredients originating from CHina, including San Francisco/Oakland, CA (60.36%), Seattle, WA (20.54%), Baltimore, MD (4.13%), and Los Angeles, CA (3.78%).

This is a new approach to analyze the risk management of feed imports, focusing on seaport of highest risk and quantity of product received. This work represents an initial step towards building a comprehensive listing of imported products introduced into the pork supply chain, and provide a roadmap to understanding risks involved in global livestock feed ingredient sourcing.

## Introduction

As an epidemic of African swine fever virus (ASFV) continues to sweep across China, the US swine industry remains in a constant state of high alert. As news of the devastating disease abroad continues to make headlines, pork producers in the US are left wondering if the safeguards practiced at established ports of entry into the US will be enough to keep ASFV from infiltrating our shores. Although once not considered a practicable means of disease transmission, the possibility of ASFV transport and introduction into the US via the importation of contaminated swine feed ingredients from China has become a topic of discussion as new scientific evidence emerges.

Specifically, physical evidence (ASFV DNA through PCR) has been detected in the Chinese feed system, specifically in raw grains and meals drying on the ground and in milling facilities and feed delivery vehicles (1). In addition, experimental evidence of ASFV survival in multiple ingredients routinely fed to pigs during a simulated 30-day trans-Atlantic journey has been published (2). Among other viruses, ASFV remained viable in certain ingredients, including three different soy products: conventional (high protein/low fat) soybean meal, organic (low protein/high fat) soybean meal and soy oil cake. Recent work by Niederwerder reported the half-life of ASFV in conventional soybean meal, organic soybean meal and soy oil cake to be 9.6, 12.9 and 12.4 days, respectively (3). In addition, survival of porcine epidemic diarrhea virus (PEDV) has been demonstrated in conventional and organic soybean meal in a simulated 37-day trans-Pacific journey, and for up to 180 days post-inoculation in conventional soybean meal following exposure to cool environmental temperatures (4,5). Furthermore, transmission of PEDV and ASFV to naïve pigs following consumption of contaminated feed has been demonstrated (6,7), and the minimum infectious dose in swine feed has been calculated for both viruses (7,8), thereby strengthening the argument that feed is a potential risk factor for the domestic and transboundary spread of certain viral pathogens.

With the understanding that soy-based products may serve as a vehicle for pathogen entry into the US, the next step is determining how much of, and more importantly, where these ingredients are entering the country. This is especially challenging considering that there are currently 328 air, land, and sea ports of entry that are overseen by US Customs and Border Protection (CBP), and that on annual basis, approximately 2.4 million metric tons of agricultural products are imported into the USA from China (2). However if for instance, it can be demonstrated that the vast majority of soy-based feed ingredients enter the US from a handful of distinct ports of entry, then the monumental task of determining where to focus limited surveillance resources becomes both manageable and effective. Therefore, the purpose of this study was to develop an analytic tool to determine the quantity of soy-based ingredients entering the US from China, and identify high-risk seaports that handle the bulk of these ingredients. This information would then research provide a simple and effective tool for risk prioritization when responding to and developing prevention protocols in response to foreign animal disease threats as they continue to emerge.

## Methods

Information on the quantity of soy-based feed ingredients and their specific ports of entry was obtained at the International Trade Commission Harmonized Tariff Schedule website (www.hs.usitc.gov), a publically available website that provides a transaction of specific trade commodities between the US and its international trading partners. Each trade commodity in the USITC database is identified by a specific 10-digit code known as the Harmonized Tariff Schedule (HTS), which is used for determining tariff classifications for all goods imported into the United States. Each commodity is classified based on the product’s name, use, and/or the material type, resulting in over 17,000 unique classification code numbers. A close review of this database identified a total of 8 HTS codes that pertain to soy-based feed ingredients, including soybeans, soybean meal, soy oil cake and soy oil.

Importing countries to be examined was limited to the 43 ASFV-positive countries currently listed on the Canadian Food Inspection Agency (CFIA) ASFVV Watch List (reference). These countries, spread across Asia, Africa, and Europe, have been determined high-risk areas for potential ASFV contamination of feed. Inclusion is based on the presence of circulating ASFV in the country. The CFIA has recently imposed stricter permitting and surveillance of raw and unprocessed grains originating from these countries that arrive at Canadian POEs (CFIA). A comprehensive list of these countries is available in *Appendix A*.

Specific queries on the 8 HTS codes and 43 countries of interest were designed on the USITC website to create a comprehensive analysis providing information on country of origin, quantity of product, year of entry, and POE into the US for each HTS code. Data was than exported into Microsoft Excel, where it could be organized and filtered into pivot tables that describe the quantity of specific product by country of origin and POE. The pivot tables display data in a manner that ranks the quantity of each product that comes into the US, its country of origin, and also its port of entry (POE). This information is further broken down into percentages to determine where the majority of high risk feed ingredients are entering the US.

A similar process was utilized to examine a five-year analysis on specific soy-based ingredients to demonstrate changes in volume and POEs into the United States. The analysis allows the viewer to easily determine how much of and where certain products of interest are entering the country. These pivot tables feature a flexible interface that allows specific products to be viewed in greater or lesser detail and ranked in comparison to other importing countries or POEs.

## Results

### Soy-based products

Eight specific 10-digit HTS codes were identified as soy-based commodities with the potential to be included in swine diets. Each code specifies pertinent details about the commodity for the purpose of tariff classification at US POEs. These HTS codes, along with their USITC database description, are provided in Table 1.

**Table 1:** Categorization of soy-based commodities arriving at US POEs of importation

|  |  |
| --- | --- |
| **HTS Code** | **Description** |
| 1201.10.0000 | FLOURS AND MEALS OF SOYBEANS, NESOI* |
| 1201.90.0005 | SOYBEAN OIL AND ITS FRACTIONS, FULLY REFINED, WASHED, BLEACHED OR DEODORIZED BUT NOT CHEMICALLY MODIFIED, NESOI |
| 1201.90.0010 | SOYBEAN OIL AND ITS FRACTIONS, ONCE-REFINED (SUBJECT TO ALKALAI OR CAUSTIC WASH BUT NOT BLEACHED OR DEODORIZED), NOT CHEMICALLY MODIFIED |
| 1201.90.0090 | SOYBEAN OILCAKE AND OTHER SOLID RESIDUES RESULTING FROM THE EXTRACTION OF SOY BEAN OIL, WHETHER OR NOT GROUND OR IN THE FORM OF PELLETS |
| 1208.10.0090 | SOYBEAN SEEDS OF A KIND USED AS OIL STOCK, WHETHER OR NOT BROKEN |
| 1507.90.4020 | SOYBEAN SEEDS OF A KIND USED FOR SOWING |
| 1507.90.4040 | SOYBEANS, CERTIFIED ORGANIC, WHETHER OR NOT BROKEN, EXCEPT SEEDS OF A KIND USED FOR SOWING OR USED AS OIL STOCK |
| 2304.00.0000 | SOYBEANS, WHETHER OR NOT BROKEN, OTHER THAN CERTIFIED ORGANIC, NESOI |
| *NESOI: Referes to “Not Elsewhere Specified or Indicated” |  |

In the year 2018, the United States imported a total of 104,707 metric tons (MT) of these commodities from nine countries included on the CFIA ASFV Watch List. The nine countries include China, Ukraine, Russia, Uganda, Taiwan, Belgium, Togo, Vietnam, and Thailand. Of this total volume, a total of 55,101 MT, or 52.6% of these soy based ingredients were imported into the US from China. Ukraine was the second largest exporter of soy-based products into the US in 2018, with 44,776 MT (42.8%) of product, and Russia being the third largest at 3,396. MT (3.2%). Each of the remaining countries on the list accounted for less than 1% of total soy-based product imported into the US. These data are presented in Table 2.

**Table 2:** Total Volume and Country of Origin of Soy-based Imports in 2018

| **Table 2: Total Volume and Country of Origin of Soy-based Imports in 2018** |  |  |
| --- | --- | --- |
| **Country of Origin** | **Sum of Year 2018 (Metric Tons)** | **% of Total** |
| China | 55,101 | 52.6% |
| Ukraine | 44,776 | 42.8% |
| Russia | 3,396 | 3.2% |
| Uganda | 990 | 0.9% |
| Taiwan | 273 | 0.3% |
| Belgium | 143 | 0.1% |
| Togo | 223 | 0.0% |
| Vietnam | 3 | 0.0% |
| Thailand | 2 | 0.0% |
| Grand Total | 104,707.0 | 100.0% |

Using data sorting filters, and the “expand” feature of the pivot table, the user can further explore the import profile of individual countries. For example, import information on China can be expanded to breakdown the total overall volume of individual soy-based ingredients as they represent the parts of the whole (Table 3). In this table, it is revealed that ground or pelletized soy-oil cake accounts for 41,998 MT, or 76.2% of all soy based products entering the US from China in 2018.

**Table 3:** Volume Analysis of Individual Soy-based Ingredients from China into US in 2018

|  |  |  |
| --- | --- | --- |
| **China** | **Sum of Year 2018 (Metric Tons)** | **% of Total** |
| SOYBEAN OILCAKE AND OTHER SOLID RESIDUES RESULTING FROM THE EXTRACTION OF SOY BEAN OIL, WHETHER OR NOT GROUND OR IN THE FORM OF PELLETS | 41,998 | 76.22% |
| SOYBEANS, CERTIFIED ORGANIC, WHETHER OR NOT BROKEN, EXCEPT SEEDS OF A KIND USED FOR SOWING OR USED AS OIL STOCK | 7,780 | 14.12% |
| SOYBEANS, WHETHER OR NOT BROKEN, OTHER THAN CERTIFIED ORGANIC, NESOI | 5,112 | 9.28% |
| SOYBEAN OIL AND ITS FRACTIONS, ONCE-REFINED (SUBJECT TO ALKALAI OR CAUSTIC WASH BUT NOT BLEACHED OR DEODORIZED), NOT CHEMICALLY MODIFIED | 103 | 0.19% |
| SOYBEAN SEEDS OF A KIND USED FOR SOWING | 62 | 0.11% |
| FLOURS AND MEALS OF SOYBEANS, NESOI | 21 | 0.04% |
| SOYBEAN SEEDS OF A KIND USED AS OIL STOCK, WHETHER OR NOT BROKEN | 18 | 0.03% |
| SOYBEAN OIL AND ITS FRACTIONS, FULLY REFINED, WASHED, BLEACHED OR DEODORIZED BUT NOT CHEMICALLY MODIFIED, NESOI | 7 | 0.01% |
| **Grand Total** | **55,101** | **100.00%** |

The next feature of these data sets is the ability of the user to reorganize the display to include individual US POEs. For example, Table 4 reveals the volume and percentage of the 55,101 MT of soy based ingredients from China that enter the US at specific POE. In this table, it is revealed that soy-based ingredients from China entered the US from 13 separate POEs in 2018. Of these POEs, a total of 4 POEs received greater than 88% of all of soy-basedingredients, including San Francisco/Oakland, CA (60.36%), Seattle, WA (20.54%), Baltimore, MD (4.13%), and Los Angeles, CA (3.78%).

**Table 4:** POE Volume Analysis of Soy-based Ingredients from China into US in 2018

|  |  |  |
| --- | --- | --- |
| **Importing Country and POE** | **Sum of Year 2018 (Metric Tons)** | **% of Total** |
| **China** | **55,101** | **100.00%** |
| San Francisco, CA | 33,261 | 60.36% |
| Seattle, WA | 11,318 | 20.54% |
| Baltimore, MD | 2,275 | 4.13% |
| Los Angeles, CA | 2,085 | 3.78% |
| Chicago, IL | 1,533 | 2.78% |
| Norfolk, VA | 1,526 | 2.77% |
| New Orleans, LA | 1,484 | 2.69% |
| Houston-Galveston, TX | 681 | 1.24% |
| Columbia-Snake, OR | 433 | 0.79% |
| New York, NY | 417 | 0.76% |
| Great Falls, MT | 60 | 0.11% |
| Savannah, GA | 18 | 0.03% |
| Minneapolis, MN | 11 | 0.02% |
| **Grand Total** | **55,100.9** | **100.00%** |

Results thus far have presented classification of soy-based ingredients at US POEs, total volumes and percentages based on country of origin, and individual volumes of specific products coming from select countries into specific POE. By compiling similar numbers over the period of 2014-2018, a five-year trend on soy-based imports can be constructed. Figure 1 summarizes the total volume in metric tons of all soy-based ingredients (8 codes) that have entered the US from China through the top four 2018 POEs, plus the 15 remaining POEs compiled as one, over the past five years.

**Figure 1:**
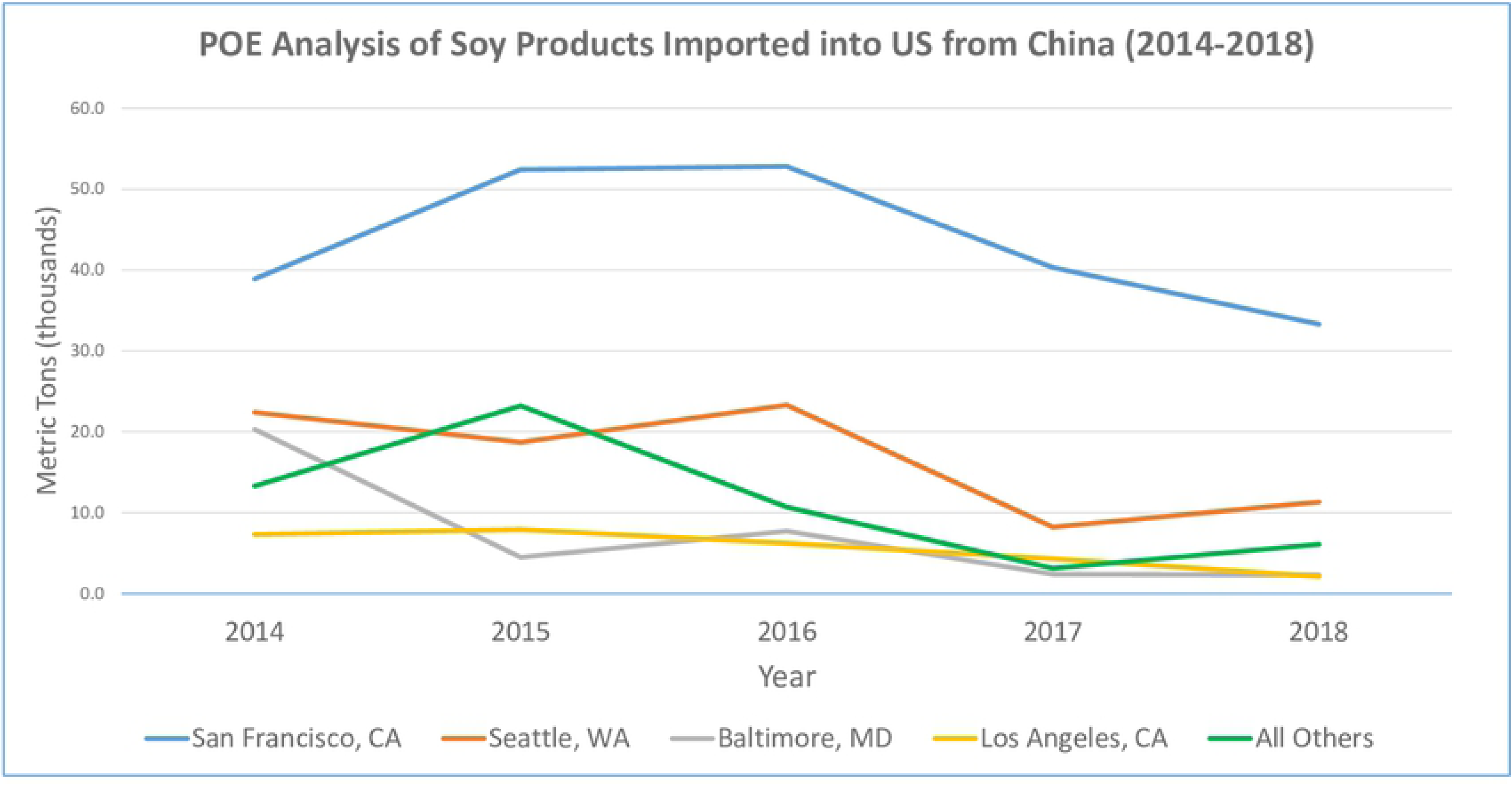

## Discussion

As the US feed supply becomes increasingly globalized, the significance and risk of foreign animal disease events as they occur is multifold, particularly when dealing with agricultural trade commodities from affected countries. Although expanding international trade allows access to diverse and competitive trade markets, the loss in direct oversight reduces consumer confidence in commodity quality control and safety. Recent scientific studies have confirmed that numerous foreign animal viruses, such as ASFV, are capable of surviving for extended time periods in select ingredients that are commonly fed to pigs and other livestock (2). Additional work has confirmed that pigs are susceptible to infection with ASFV through the consumption of contaminated plant-based feed (7). Therefore, it is imperative that swine feed ingredients imported into the US from ASFV positive countries be treated with increased scrutiny and caution.

Knowledge of this novel risk factor seemingly presents an immeasurable challenge for US Customs and Border Protection (CBP) due to the sheer volume of imported product. As state previously, the US imports approximately 2.4 million metric tons of agricultural products such as meat, grain, vitamins, minerals, and amino acids from China on an annual basis (2). Across the 328 air, land, and sea ports of entry that CBP oversees, it is an additional challenge to decide where, and over which particular products, the application of limited resources and oversight will be most effective. Without unlimited resources, rational decisions must be made on where to concentrate additional surveillance efforts, with a focus on areas where risk of disease entry are highest. This study represents an attempt to quantify the amount and identify the place of entry for high-risk feed ingredients originating from ASFV-positive countries. Understanding these metrics is the first step to risk prioritization efforts, and is necessary for appropriate response actions to be taken.

Another novel feature of this study is the ability to determine specific ports of entry for various products of interest. This information, combined with historic data to support the relatively stable flow of particular products through specific ports over time, is critical to understanding the mostly likely ASFV entry points. While it may come as no surprise that the majority of products originating from China are received at US POEs along the Pacific coast, having firm numbers to compare and contrast helps to justify where additional ASFV-entry mitigation resources will be most cost-effective.

With the generation of new knowledge on viral half-life in feed, the application of a “Responsible Imports” approach has been adapted across the US industry (2). Responsible Imports, a science-based protocol to safely introduce essential feed ingredients from high-risk countries, is based on the following principles:

**Necessity**: is importation of the ingredient an absolute necessity?

**Alternatives:** can the ingredient be obtained locally or from a country free from foreign animal diseases?

**Virus:** which virus is causing the concern?

**Viral half-life:** is there published information on the half-life of the virus in the designated ingredient?

**Transport time:** what is the projected time for delivery of the ingredient from the source to its destination?

**Viral load:** are there safe products that can be added to the ingredient to reduce viral load during transport?

**Storage period:** is there published information on storage time and temperature that will reduce residual virus from the ingredient prior to use?

Therefore, as production companies across the US develop storage facilities for incoming products, a new way of thinking is taking shape: one based on “feed quarantine” that brings together information across several fronts including feed science, microbiology and oceanic transport logistics to understand how to minimize risk. This approach is intriguing as it is non-regulatory in nature, and does not negatively impact trade.

It should be noted that the results of this study are not without limitations. As shown in table 2, 55.1 thousand metric tons of soy-based feed ingredients entered the US from China in 2018. It is important to consider that this number represents, along with all numbers presented in the study, the total amount of product cleared by US Customs at US POEs. USITC defines these products as “imports for consumption,” intended for use and distribution across all industries and markets, and does not provide any further information on final product destination or intended use. It is not possible therefore, using only the methods presented in this study, to determine how much of a particular product ultimately ends up in the domestic swine supply chain.

However, this gap in knowledge does not negate the potential risk of contaminated feed reaching the US swine population. Modern swine diets contain hundreds of ingredients, all of which can be mixed and matched for consumption by pigs. Additionally, many of these rations are prepared at individual feed mills that then distribute the complete feed product to numerous farms across a region. While a great deal of feed material may be sourced locally, one 2018 analysis found that inventory at a Midwestern swine farm included ingredients from 12 countries in North America, Asia, and Europe (8). It is therefore entirely possible that ingredients might make their way from ASFV-positive countries to U.S. farms. And, crucially, it might not be the ingredients themselves that matter the most: the trucks and packaging that carry them may also be capable of spreading disease.

While information on soy-based products is immediately applicable in developing response strategies for ASFV, there is also a wider utility and potential that is revealed from this analytical process. Increased globalization has also brought a greater number of adverse health events that have been attributed to contaminated or infectious food commodities imported from abroad and consumed by both animals and humans. The hazards that foreign food commodities have the potential to possess include products that are adulterated either intentionally or unintentionally, foreign infectious diseases, and dangerous contaminates. This same analysis and organizational method can be applied to nearly any foreign trade commodity of interest. In fact, the USITC website contains over 17,000 unique trade commodities, organized by searchable terms that can, by using this data analysis process, be displayed in an easy to understand format. Applying this information to risk prioritizations plans can have broad reaching implications for the development of both human and animal food safety protocols.

In closing, this analytical tool provides an opportunity to gather information that is important in developing science-based plans to safely import essential ingredients that we cannot manufacture in the US, along with the selective exclusion of ingredients of limited value and present significant risk. It is hoped that these efforts will continue to stimulate communication and collaboration between the feed and livestock industries, resulting in further research into the emerging concept of “global feed biosecurity”. Ideally, current and future information regarding the risk of pathogen spread in feed will enhance the accuracy of risk assessments, drive the continual development of efficacious feed-based mitigation strategies and ultimately, bring the health status in the country of origin into the forefront of philosophies regarding the global trade of feed ingredients.

## Appendix A CFIA Watch List Countries

**Appendix A: CFIA Watch List Countries**
|  |
| --- |
| Belgium |
| Benin |
| Burkina Faso |
| Bulgaria |
| Burundi |
| Cabo Verde |
| Cameroon |
| Central African Republic |
| Chad |
| China |
| Congo |
| Cote D'Ivoire |
| Czech Republic |
| Estonia |
| Gambia |
| Ghana |
| Guinea-Bissau |
| Hungary |
| Italy |
| Kenya |
| Latvia |
| Lithuania |
| Madagascar |
| Malaw |
| Moldova |
| Mongolia |
| Mozambique |
| Namibia |
| Nigeria |
| Poland |
| Romania |
| Russia |
| Rwanda |
| Senegal |
| Sierra Leone |
| South Africa |
| Tanzania |
| Togo |
| Uganda |
| Ukraine |
| Vietnam |
| Zambia |
| Zimbabwe |

## References

1. Proceedings of the 1st International Symposium on the Prevention of ASFV, the Chinese Pig Producers Association, Henan, China, February 22, 2019.

2. Patterson, G., Niederwerder, M. C., & Dee, S. A. (2019). Risks to animal health associated with imported feed ingredients. Journal of the American Veterinary Medical Association, 254(7), 790–791. https://doi.org/10.2460/javma.254.7.790.

3. Dee, S. A., Bauermann, F. V., Niederwerder, M. C., Singrey, A., Clement, T., de Lima, M., … Diel, D. G. (2018). Survival of viral pathogens in animal feed ingredients under transboundary shipping models. PLOS ONE, 13(3), e0194509. https://doi.org/10.1371/journal.pone.0194509

4. Stoian AMM, Zimmerman J, Ji J, Hefley TJ, Dee S, Diel DG, Rowland RRR, Niederwerder MC. Half-Life of African Swine Fever Virus in Shipped Feed. Emerg Infect Dis. 2019 Dec 17;25(12). doi: 10.3201/eid2512.191002.

5. Dee S, Neill C, Singrey A, Clement T, Cochrane R, Jones C, Patterson G, Spronk G, Christopher-Hennings J, Nelson E. Modeling the transboundary risk of feed ingredients contaminated with porcine epidemic diarrhea virus. BMC Vet Research. 2016 Mar 12;12:51. doi: 10.1186/s12917-016-0674-z.

6. Dee S, Neill C, Clement T, Singrey A, Christopher-Hennings J, Nelson E. An evaluation of porcine epidemic diarrhea virus survival in individual feed ingredients in the presence or absence of a liquid antimicrobial. Porcine Health Manag 2015 Jul 9;1:9. doi: 10.1186/s40813-015-0003-0. eCollection 2015.

7. Dee S, Clement T, Schelkopf A, Nerem J, Knudsen D, Hennings J, Nelson E. An evaluation of contaminated complete feed as a vehicle for porcine epidemic diarrhea virus infection of naïve pigs following consumption via natural feeding behavior: Proof of concept. BMC Vet Res. 2014;10:176.

8. Niederwerder MC, Stoian AMM, Rowland RRR, Dritz SS, Petrovan V, Constance LA, Gebhardt JT, Olcha M, Jones CK, Woodworth JC, Fang Y, Liang J, Hefley TJ. Infectious Dose of African Swine Fever Virus When Consumed Naturally in Liquid or Feed. Emerg Inf Dis. 2019 May;25(5):891–897. doi: 10.3201/eid2505.181495. Epub 2019 May 17.

9. Schumacher LL, Woodworth JC, Jones CK, Chen Q, Zhang J, Gauger PC, Stark CR, Main RG, Hesse RA, Tokach MD, Dritz SS. Evaluation of the minimum infectious dose of porcine epidemic diarrhea virus in virus-inoculated feed. Am J Vet Res 2016 Oct;77(10):1108–13. doi: 10.2460/ajvr.77.10.1108.

